# Severe Myocardial Inflammation and Necrosis in Juvenile Mice Compared with Adult Mice with Coxsackievirus B3 Myocarditis

**DOI:** 10.64898/2026.08.20.746108

**Authors:** Jacob C Ricci, Logan P Macomb, Emily R Whelan, Katherine Gegoutchadze, Cormac J Davis, Kyra G Ritter, Priya Tomerlin, Ashley A Darakjian, Nick A Farahani, Lauren M Parrow, Danielle J Beetler, Max W Strandes, Damian N Di Florio, Sami Khatib, Jude Elsaygh, Leslie T Cooper, Jack F Price, DeLisa Fairweather, Dipankar Gupta, Katelyn A Bruno

## Abstract

**Background:** Viral myocarditis presents a significant burden of disease, particularly among children and young adults. However, clinical guidelines and treatment strategies for pediatric patients are derived from those for adult patients due to a lack of pediatric data. Current animal models of viral myocarditis use adult mice, so conclusions from these models cannot necessarily be extrapolated to the pediatric population. We sought to develop a juvenile mouse model of myocarditis to examine differences between these two distinct clinical populations.

**Methods:** Male and female BALB/c 3-4-week-old “juvenile” and 8-week-old “adult” mice were infected intraperitoneally with 10^3^ PFU of heart-passaged coxsackievirus B3. Sera was used to evaluate testosterone and estradiol levels. Cardiac histological evaluations included overall inflammation, fibrosis, and specific cell-type infiltration. RNA was extracted from cardiac tissue and evaluated for changes in gene expression of cell-type markers, complement components, and NLRP3 inflammasome components.

**Results:** Juvenile mice exhibited more severe inflammation than adult mice but no sex differences in overall inflammation. Juvenile mice demonstrated iancreased infiltration of CD11b+ cells, F4/80+ cells, and CD3+ T-cells vs. adults. Inflammasome genes NLRP3 and caspase-1 were significantly increased in juvenile compared with adult myocarditis.

**Conclusions:** This paper is the first to describe a juvenile mouse model of coxsackievirus B3 myocarditis and provides a direct comparison to a translational adulat mouse model. Juvenile mice had greater cardiac inflammation than adults. This model replicates clinical populations and provides a valuable tool to study age as a factor in the pathogenesis of myocarditis.

## BACKGROUND

Myocarditis, defined as inflammation of the myocardium, remains an important cause of sudden cardiac death among children and young adults.^1,2^ Myocarditis is characterized by immune cell infiltration into the heart and is primarily caused by viral infections, including coxsackievirus B3 (CVB3), among patients in North America.^3–8^ Within this patient population, roughly 30% of patients progress to dilated cardiomyopathy (DCM) and chronic heart failure (HF).^9,10^ Currently, there are no disease-specific medications for myocarditis, and guidelines recommend treatment of symptomatic heart failure when indicated.^11–13^ The spectrum of presentation and outcomes of myocarditis in both children and adults varies significantly.^14^ Though it is well established that myocarditis is particularly deadly in newborns and infants, specific studies focusing on age remain scarce.^15–17^ Children tend to present with a more severe or fulminant myocarditis, progress rapidly, and develop persistent cardiac dysfunction as well as an increased risk of death or need for transplantation.^16–18^ Despite differences in presentation, clinical progression, and outcomes of myocarditis across age and sex, limited research has specifically examined these factors in juvenile and adult animal models.

Among adults, males tend to develop myocarditis more often, with a ratio upwards of 3.5:1.^19,20^ Moreover, studies have shown that men are twice as likely to present with evidence of cardiac fibrosis on magnetic resonance imaging, progress from myocarditis to DCM more frequently, and have lower transplant-free recovery.^9,21^ However, the mechanisms underlying this sex difference remains poorly understood.

Animal models have been critical for understanding the pathogenesis of myocarditis and its associated sex differences.^22,23^ From these models, it has been shown that pathogenic inflammation is associated with upregulation of specific innate and adaptive pathways. Particularly, the NOD-, LRR- and pyrin domain-containing protein 3 (NLRP3) inflammasome activation via Toll Like Receptor (TLR) signaling is activated in response to coxsackievirus infection and is critical to the development of myocardial inflammation.^24,25^ Furthermore, this pathway has been shown to have greater activation in males compared with females at multiple points within the pathway.^25,26^ However, these experiments have focused on adult mice. To this point, comparison of responses in young mice in other established mouse models of myocarditis have not been conducted. Therefore, the goal of this work was to develop and describe a new animal model of juvenile myocarditis.

## METHODS

### Animal Care Ethics Statement

Mice were used in strict accordance with the recommendations in the Guide for the Care and Use of Laboratory Animals from the National Institutes of Health. Mice were maintained under pathogen-free conditions in the animal facilities at Mayo Clinic, Florida. Approval was obtained from the Animal Care and Use Committee at Mayo Clinic Florida for all procedures (IACUC number: A00004548-19-R22). Mice were sacrificed according to the Guide for the Care and Use of Laboratory Animals from the National Institutes of Health.

### Juvenile and Adult CVB3-Induced Myocarditis Models

Male and female BALB/c 3-4-week-old “juvenile” and 8-week-old “adult” mice were obtained from Jackson Laboratories (Bar Harbor, ME, #000651). Mice were housed in pathogen-free conditions at the Mayo Clinic animal facility in Florida. Due to the inability to purchase mice under 3 weeks of age and inconsistencies in the exact ages of mice obtained from Jackson Labs, puberty checks were performed to determine puberty status.^27,28^ Females were deemed prepubescent if the vaginal opening was not seen. Males were considered prepubescent if there was no preputial separation. Mice were inoculated with sterile phosphate-buffered saline (PBS) or 10^3^ plaque-forming units (PFU) of heart-passaged stock of CVB3 via intraperitoneal (ip) injection on day 0. Acute myocarditis was examined on day 10 post-infection (pi) as previously described for adults.^22,29^ The same time course was used for the juvenile model. CVB3 (Nancy Strain) was obtained from the American Type Culture Collection (ATCC, Manassas, VA, VR-30), grown in Vero cells (ATCC, Manassas, VA, CCL-81), and passaged through the heart, as previously described.^22,30^

### Histology

#### Myocarditis

Hearts were cut longitudinally and submerged in 10% phosphate-buffered formalin to fix tissue for 24-72 hours. Hearts were embedded in paraffin before sectioning. Five-micron sections were cut and stained with hematoxylin and eosin (H&E) to detect inflammation. Myocarditis was assessed as the percentage of the heart with inflammation compared to the overall size of the heart section based on visual analysis using a microscope eyepiece grid, as previously described.^31–35^ Sections were scored by two individuals blinded to the groups.

#### Fibrosis

Additional heart sections (ten-micron) were cut and stained using picrosirius red to detect fibrosis. Picrosirius red stain was prepared by dissolving 0.5 g of Sirius Red F3B (Thermo Fisher Scientific) in 500 mL of saturated aqueous picric acid (Thermo Fisher Scientific). After deparaffinizing and rehydrating, slides were incubated in picrosirius red solution for 1 hour. Resulting slides were scanned using an Aperio ScanScope CS2 (Leica, Wetzlar, Germany). Tissue annotations were created using QuPath (Version 0.6.0). All annotations were reduced by 30 μm to exclude the pericardium and subsequently adjusted to include only ventricular tissue. The percent fibrosis was determined using a pixel thresholder (Threshold value: 0.3) to obtain a collagen-positive area, which was divided by the total area of the annotated tissue.

#### Immunohistochemistry

Five-micron sections of hearts were stained for CD45 (Biolegend, San Diego, CA, USA, 103102, 1:200, rat), CD11b (Abcam, Cambridge, United Kingdom, ab133357, 1:3000, rabbit), F4/80 (BioRad, Hercules, CA, USA, MCA497G, 1:250, rat), and CD3 (Abcam, ab16669, 1:200, rabbit). Secondary antibodies from Envision+ anti-rabbit (K4006) and rat-on-rodent (RT517) kits (Biocare, Pacheco, CA, USA) were used for rabbit and rat antibodies, respectively. Visualization was achieved using 3,3’-diaminobenzidine (DAB)+ as the substrate-chromogen. Stained slides were scanned using an Aperio CS2 scanner (Leica, Wetzlar, Germany). Heart tissue was annotated using a pixel thresholder in QuPath and subsequently adjusted only to include ventricular tissue. Stain positivity (% Positive) was determined for CD45 using a pixel thresholder (resolution: 0.26 μm/pixel; threshold value: 0.15). CD11b, F4/80, and CD3 were detected using QuPath’s positive cell detection algorithm. The numeric variables for each marker were as follows: CD11b (Requested Pixel Size: 0.25 μm, Background Radius: 10.0 μm, Median Filter Radius: 0.0 μm, Sigma: 1.5 μm, Minimum Area: 2.0 μm^2^, Maximum Area: 400.0 μm^2^, Threshold: 0.07, Max Background Intensity: 2.0, Positive Threshold: 0.2); F4/80 (Requested Pixel Size: 0.5 μm, Background Radius: 5.0 μm, Median Filter Radius: 0.0 μm, Sigma: 1.5 μm, Minimum Area: 5.0 μm^2^, Maximum Area: 400.0 μm^2^, Threshold: 0.08, Max Background Intensity: 2.0, Positive Threshold: 0.4); CD3 (Requested Pixel Size: 0.25 μm, Background Radius: 10.0 μm, Median Filter Radius: 0.0 μm, Sigma: 1.5 μm, Minimum Area: 5.0 μm^2^, Maximum Area: 4000.0 μm^2^, Threshold: 0.07, Max Background Intensity: 2.0, Positive Threshold: 0.4). Percent positivity was determined as positive # cells/total # cells.

### RNA Isolation from Heart Tissue

On day 10 pi, half of the heart was collected and flash-frozen before storage at −80° C for RNA isolation. Hearts were lysed and homogenized in RLT buffer with 0.5% DX buffer using a 7mm stainless steel bead and Tissuelyser LT (Qiagen, Germantown, MD). RNA was then isolated from the homogenate using the QIAcube RNase Easy Fibrous Mini Kit, which includes DNase and proteinase K digestion (Qiagen, Germantown, MD). RNA was eluted into 30 µL. RNA quantification was determined in µg/µL using the NanoDrop OneC UV-Vis Spectrophotometer (Thermo fisher Scientific, Waltham, MA).

#### qRT-PCR Method

Total RNA from hearts was assessed by quantitative real-time (qRT) PCR using the ABI 7000 Taqman system or ABI QuantStudio 3 (Applied Biosystems, Foster City, CA). RNA was first converted to cDNA using a high-capacity reverse transcriptase kit (Applied Biosystems, Foster City, CA) as previously described.^36,37^ Expression was calculated as Fold Change (FC) normalized to the housekeeping gene hypoxanthine phosphoribosyltransferase 1 (HPRT). All the following primers were purchased from Thermo Fisher Scientific (Waltham, MA): CD45 (Assay ID: Mm01293577_m1); CD11b (Mm00434455_m1); F4/80 (Mm00802529_m1); CD3 (Mm01179194_m1); CD4 (Mm00442754_m1); CD8 (Mm01182107_g1); C3 (Mm01232779_m1); C4 (Mm00437893_g1); C3aR1 (Mm62620006_s1); C5aR1 (Mm00500292_s1); TLR2 (Mm00442346_m1); TLR4 (Mm00445273_m1); TLR7 (Mm04933178_g1); NLRP3 (Mm00840904_m1); Casp1 (Mm00438023_m1); Pro-IL-1β (Mm00434228_m1).

Expression was calculated using the ΔΔCt method. Briefly, each gene was analyzed by subtracting the cycle threshold (Ct) of the gene of interest from that of the housekeeping gene to determine ΔCt. The ΔCt values for 8-week-old uninfected females were averaged to give the reference ΔCt (average). This average value was then subtracted from each sample ΔCt value to provide the ΔΔCt. Fold change was calculated with the equation FC = 2^-ΔΔCt^.

#### CVB3 VP1

Probe sets to detect CVB3 VP1 were developed as previously described and obtained from Integrated DNA Technologies (Coralville, IA).^38^ Probe sets are as follows: CVB3 forward, 5′-CCCTGAATGCGGCTAATCC-3′; CVB3 reverse, 5′- ATTGTCACCATAAGCAGCCA-3′; CVB3 probe, 5′-FAM-TGCAGCGGAACCG-TAMRA-3′.

### ELISAs

Blood was collected from each mouse via cardiac puncture. Samples were allowed to clot at room temperature for at least 15 minutes before centrifugation at 1500 g for 15 minutes at 4°C. Sera was then removed via aspiration. Enzyme-linked immunosorbent assays (ELISAs) were used to determine testosterone and estradiol levels in sera for each sample. Frozen sera aliquots were allowed to come to room temperature before use in ELISA protocols. Testosterone levels were measured using the Testosterone Parameter Assay Kit (R&D Systems, Catalog #: KGE010). Estradiol levels were measured using the Estradiol Parameter Assay Kit (R&D Systems, Catalog #: KGE014). For both, absorbance was used to calculate concentration relative to a standard curve. The assay ranges were 0.041 – 10 ng/mL and 12.3 – 3000 pg/mL for testosterone and estradiol, respectively.

### Statistical Analysis

Experimental age groups were analyzed both when combined and split by sex. Outliers were identified using the ROUT method with Q = 0.5%. Normally distributed data comparing two groups were analyzed using a two-tailed Welch’s *t*-test. A two-way ANOVA with age and disease status as factors was used to investigate the effects of both age and infection. Tukey’s post hoc test was used to control for multiple comparisons. Data are expressed as scatter plots and mean ± standard error of the mean (SEM). P-values displayed are the result of a post-hoc test. All statistical calculations were performed in GraphPad Prism (Version 10.5.0 for Windows, GraphPad Software, Boston, Massachusetts, USA, www.graphpad.com), and values of p < 0.05 were considered statistically significant.

## RESULTS

### Juvenile mice exhibit greater myocardial inflammation, calcification, and fibrosis

Overall, infected juvenile mice demonstrated greater myocardial inflammation compared with infected adult mice (p < 0.0001) **(Figure 1a)**. Moreover, in both female **(Figure 1b)** and male **(Figure 1c)** mice, myocardial inflammation was greater in juvenile than adult mice (p = 0.0049 and p = 0.0157, respectively). However, juvenile mice did not show sex differences in overall percent myocardial inflammation (p = 0.89) **(Figure 1d)**. As expected, adult males had greater inflammation than females (p = 0.0381) **(Figure 1e)**. Visual assessment of histology revealed that both juvenile females and males exhibited myocardial calcification, which was not observed in adults. There was also no difference in CVB3 VP1 viral gene expression between juvenile and adult myocarditis (p = 0.29) **(Figure 2a)** or by sex (p = 0.69, p = 0.35, respectively) **(Figure 2b and 2c)**. Uninfected controls had no detectable VP1 in the heart (data not shown). Analysis of collagen levels using picrosirius red showed infected juvenile mice with myocarditis had significantly greater collagen deposition than adult mice with myocarditis (p = 0.0100) **(Figure 3a)**. However, there were no observed differences when this analysis was separated by sex **(Figure 3b and 3c)**.

**Figure 1.**
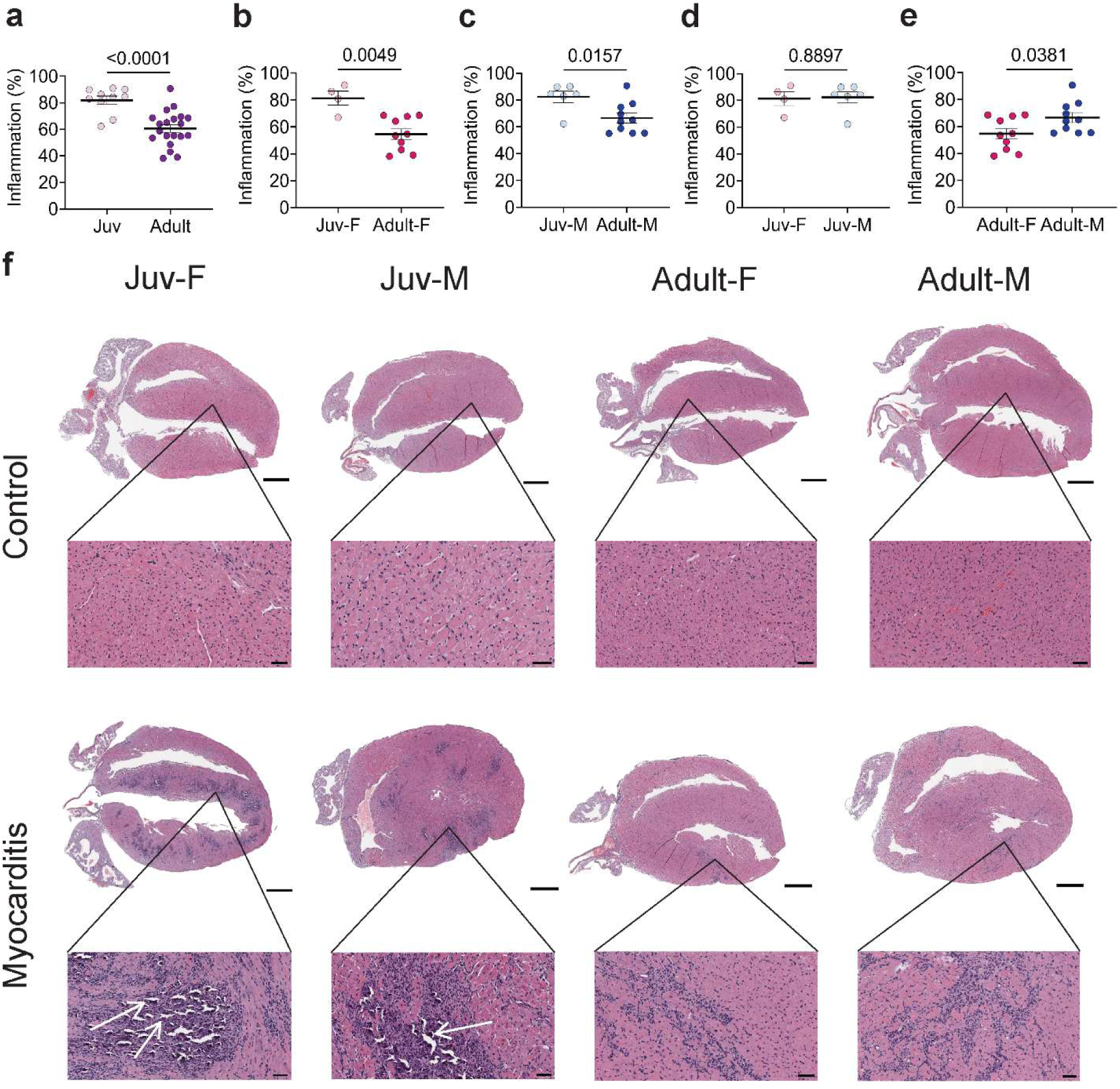
Juvenile mice with myocarditis exhibit greater cardiac inflammation than adult mice, but no sex differences. Juvenile and adult female and male mice were injected intraperitoneally with 10^3^ PFU of CVB3 and harvested at day 10 post-infection (pi). **(a)** Overall inflammation differences between infected juvenile (light purple) and adult (purple) mice were compared. Age effects on myocarditis were analyzed in **(b)** females (juvenile in pink, adult in red) and **(c)** males (juvenile in light blue, adult in dark blue). Sex differences in myocarditis were analyzed between **(d)** juvenile groups and **(e)** adult groups. Myocarditis was assessed as a percentage (%) of inflammation in histology sections using a microscope grid with hematoxylin and eosin (H&E) staining. **(f)** Representative H&E images for all groups at day 10 pi were selected. White arrows identify areas of calcification. Scale bars for whole tissue sections are 800 μm, while scale bars on callout images are 50 μm. Data shown as scatter plots and mean ± SEM using **(a-e)** two-tailed Welch’s *t*-test for analyses. n = 11 for juvenile group (4 female, 7 male). n = 20 for adult group (10 female, 10 male).

**Figure 2.**
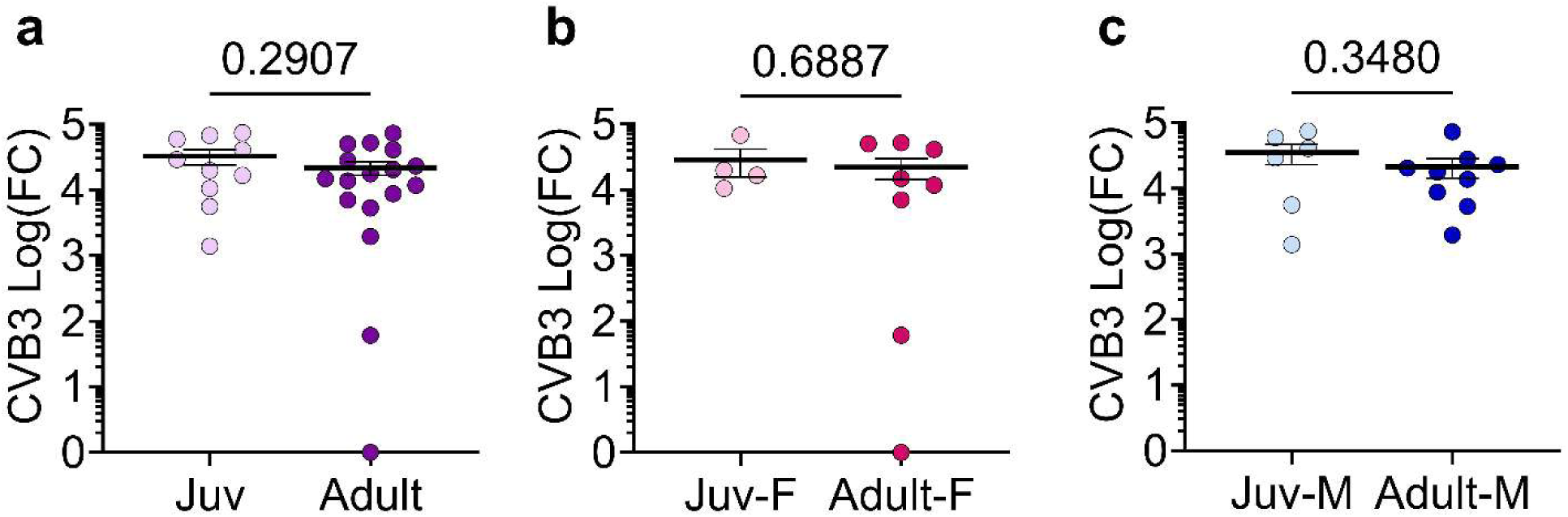
Mice with myocarditis showed no differences in cardiac viral levels regardless of sex or age group. Juvenile and adult female and male mice were injected intraperitoneally with 10^3^ PFU of CVB3 and harvested at day 10 pi. RNA was isolated and used to produce cDNA. Log fold change (FC) expression of CVB3 capsid protein VP1 in cardiac tissue was determined utilizing qRT-PCR compared to the housekeeping gene hypoxanthine phosphoribosyltransferase (HPRT). **(a)** Analysis of VP1 expression was conducted between infected juvenile (light purple) and adult (light purple) mice. An additional comparison of expression in sex-separated juvenile and adult **(b)** female (juvenile in pink, adult in red) and **(c)** male (juvenile in light blue, adult in dark blue) groups. Data shown as scatter plots and mean ± SEM utilizing two-tailed Welch’s *t*-test for analyses. n = 11 for juvenile group (4 female, 7 male). n = 20 for adult group (10 female, 10 male).

**Figure 3.**
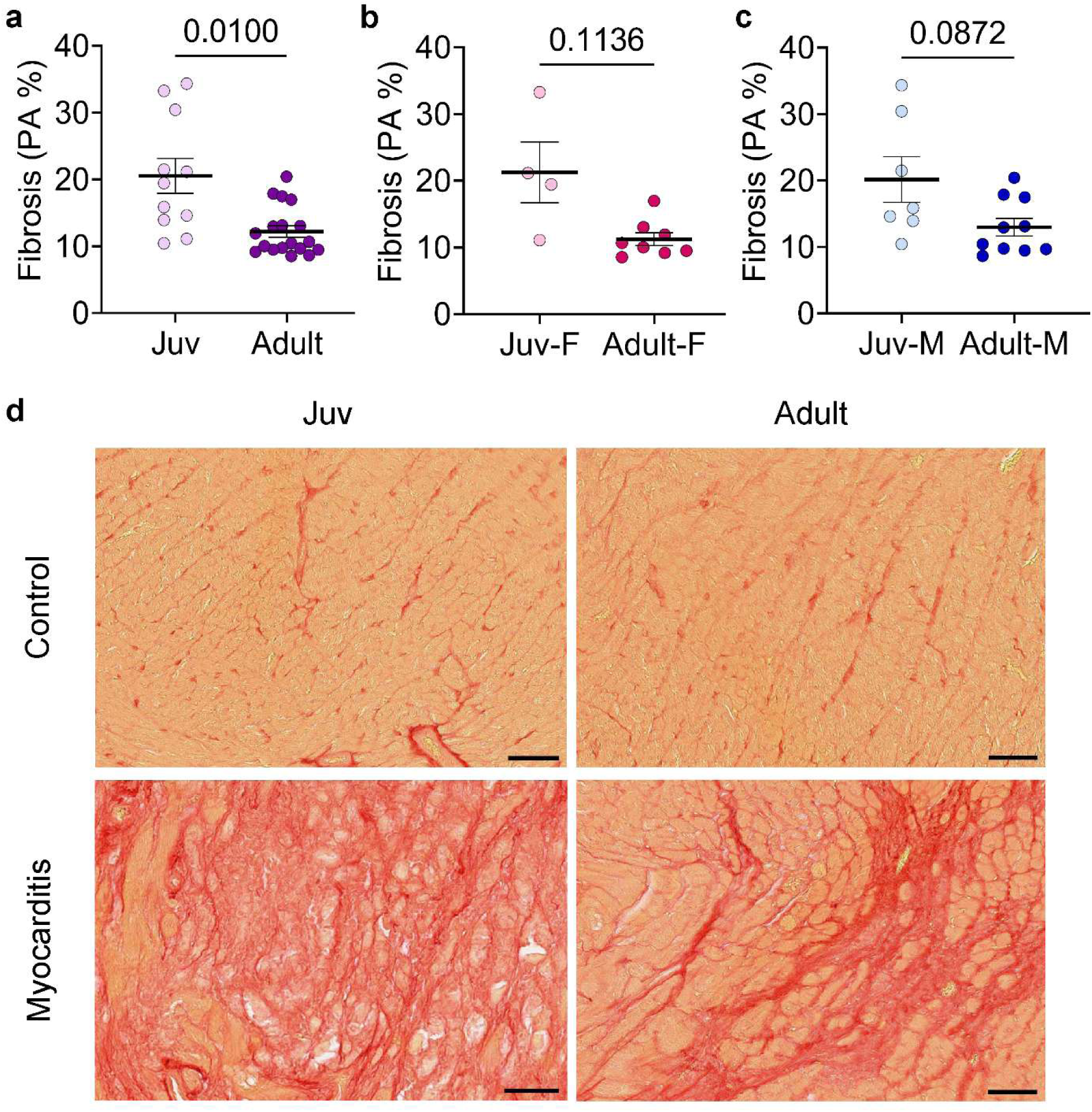
Juvenile mice with myocarditis show greater collagen content in response to viral infection than adults with myocarditis. Juvenile and adult female and male mice were injected intraperitoneally with 10^3^ PFU of CVB3, and heart sections were evaluated for collagen content via picrosirius red staining at day 10 pi. Percent positive area was determined using a pixel thresholder on QuPath. **(a)** Comparison of percent positive area was conducted between juvenile (light purple) and adult (purple) mice, regardless of sex. Further sex-separated analyses were performed between juvenile and adult **(b)** female (juvenile in pink, adult in red) and **(c)** male (juvenile in light blue, adult in dark blue) mice. **(d)** Representative images of collagen deposition. Data are presented as scatter plots and mean ± SEM utilizing two-tailed Welch’s *t*-test for analyses. n = 11 for juvenile group (4 female, 7 male). n = 20 for adult group (10 female, 10 male). All scale bars are equal to 50μm. PA % - percent positive area.

### Juvenile mice show increased cell-type specific infiltration compared with adult mice during myocarditis

Overall, supporting the finding of increased inflammation in juvenile mice, compared with adult groups, juvenile mice with myocarditis had increased CD45+ area (p = 0.0051) **(Figure 4a)**, CD11b+ cells (p = 0.0048) **(Figure 4b)**, F4/80+ cells (p = 0.0374) **(Figure 4c)** and CD3+ cells (p = 0.0043) **(Figure 4d)**. Positive area was utilized for CD45 because cell detection was not possible given the prevalence of positive staining. Analyzing by sex, juvenile males with myocarditis had increased CD45+ area (p = 0.0318) and CD11b+ cellular infiltration (p = 0.0196) into the myocardium in response to viral infection compared with adult males with myocarditis **(Figure 4e and 4f),** while juvenile females and males with myocarditis had increased CD3+ cellular infiltration compared with adult females with myocarditis (p = 0.0332, p = 0.0271, respectively) **(Figure 4h)**. Neither juvenile males nor females showed significant changes in F4/80+ cellular infiltration when compared with their sex-matched adult groups **(Figure 4g)**.

**Figure 4.**
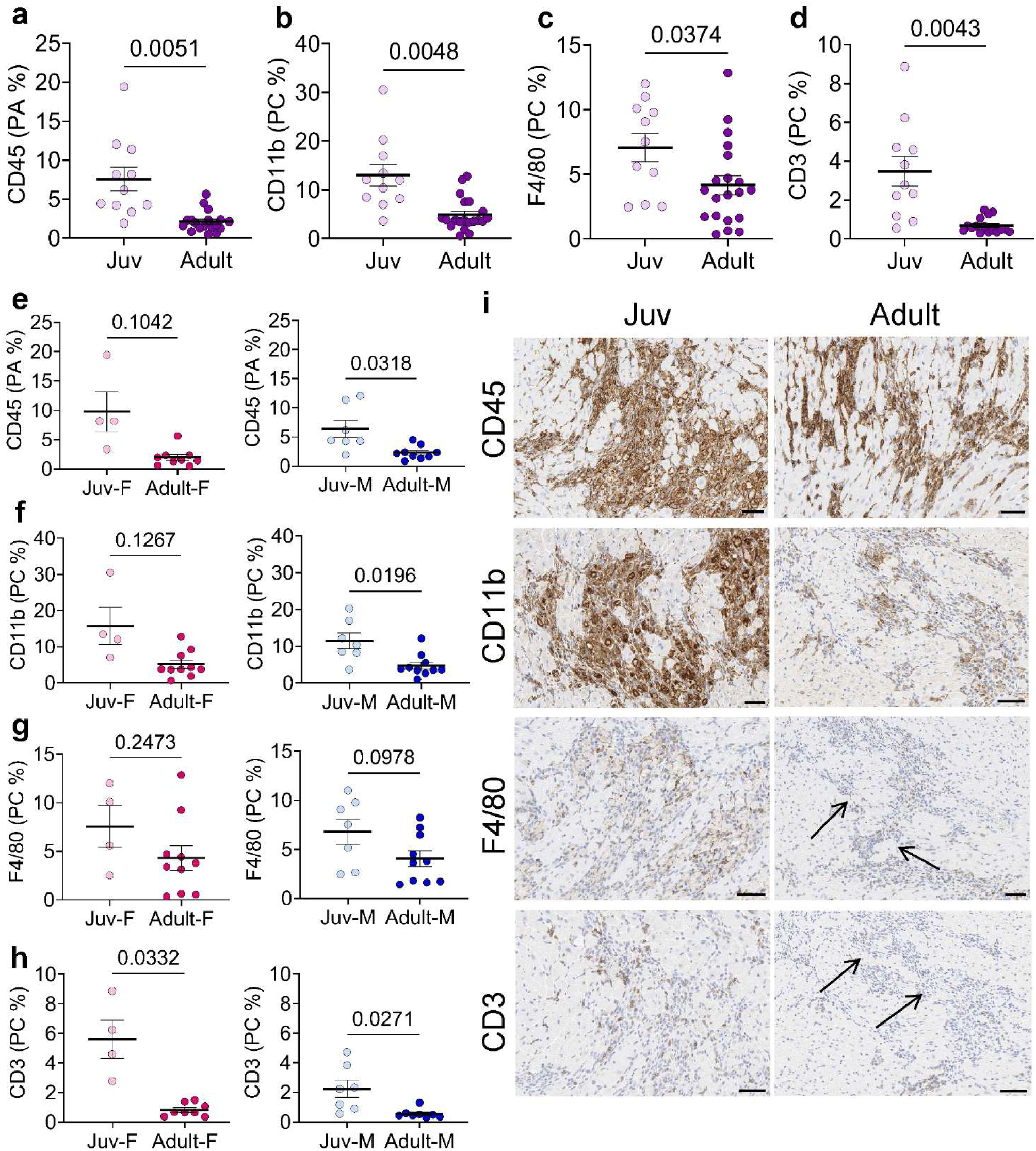
Juvenile females and males with myocarditis show differential cellular composition of immune infiltrates compared with adults. Juvenile and adult female and male mice were injected intraperitoneally with 10^3^ PFU of CVB3 and harvested atday 10 pi. Heart sections were evaluated via immunohistochemistry (IHC) staining for the targets CD45, CD11b, F4/80, and CD3. Comparisons were conducted between juvenile (light purple) and adult (purple) mice with myocarditis **(a-d)**. Further sex-separated analyses were conducted between juvenile and adult females (juvenile in pink, adult in red) and males (juvenile in light blue, adult in dark blue) for all targets **(e-h)**. **(i)** Representative images of mice with myocarditis for each target are shown. For CD45 **(a,e)**, the percent area DAB positive was used due to inability of cell detection algorithms to separate cells. For CD11b **(b,f)**, F4/80 **(c,g)**, and CD3 **(d,h)**, the percentage of DAB- positive cells was calculated. Data are presented as scatter plots and mean ± SEM utilizing two-tailed Welch’s *t*-test for analyses. n = 11 for juvenile group (4 female, 7 male). n = 20 for adult group (10 female, 10 male). All scale bars are equal to 50 μm. PA % - positive area percentage. PC % - positive cell percentage.

### Juvenile mice with myocarditis exhibit differential expression of innate and adaptive immune cell markers compared with adult mice with myocarditis

Next, we analyzed the gene expression levels of major immune cell markers within the myocardium. Compared with adult mice with myocarditis, juvenile mice with myocarditis had no changes in expression of CD11b or F4/80 **(Figure 5a and Figure 5b)**. No differences in expression between juvenile and adult mice with myocarditis were seen for CD3 (p = 0.1691), CD4 (p = 0.1355), and CD8 (p = 0.4955) (**Figure 5c, 5d, and 5e**). However, compared with adult females with myocarditis, juvenile females with myocarditis had increased expression of CD3 (p = 0.0182) **(Figure 5c)** and CD4 (p = 0.0094) **(Figure 5d).** Neither juvenile sex showed increased CD8 expression compared with adults **(Figure 5e)**. Juvenile mice had significantly greater expression of the chemokine receptor markers CX3CR1 (p = 0.0005) and CCR2 (p = 0.0064) (**Figure 6a and Figure 6b**). Both juvenile females and males with myocarditis had increased expression of CX3CR1 compared with adult mice with myocarditis (p = 0.0182 and p = 0.0101, respectively) **(Figure 6a),** while only juvenile females with myocarditis had increased CCR2 expression compared with adult females with myocarditis (p = 0.0018) **(Figure 6b)**.

**Figure 5.**
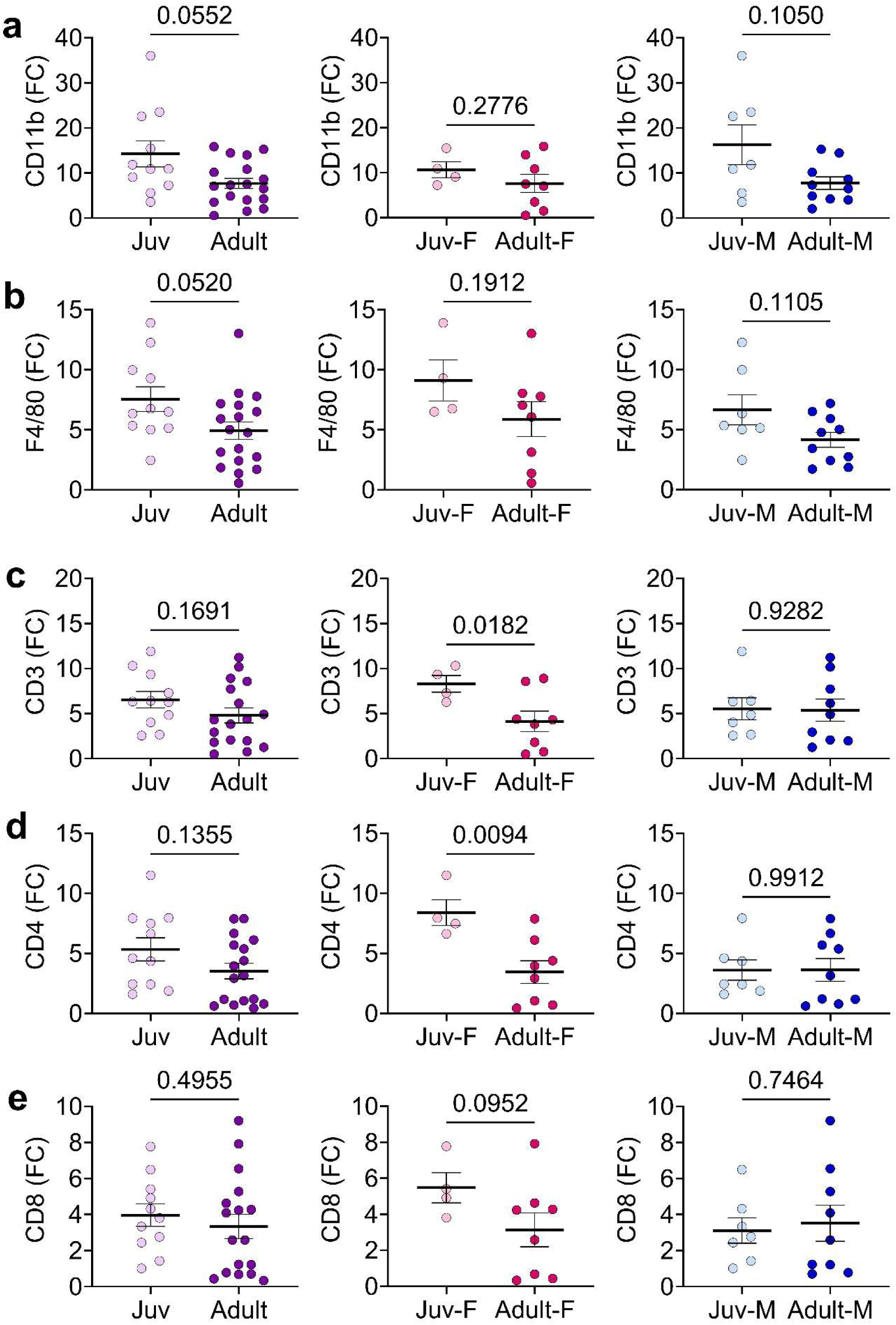
Juvenile female and male mice with myocarditis show differential increases in innate cell marker expression compared with adults. Juvenile and adult female and male mice were injected intraperitoneally with 10^3^ PFU of CVB3 and harvested at day 10 pi. RNA was isolated and used to produce cDNA. Combined groups of juvenile (light purple) and adult (purple) with myocarditis were analyzed. Further sex-separated analyses were conducted between juvenile and adult females (juvenile in pink, adult in red) and males (juvenile in light blue, adult in dark blue) for all targets. Fold change (FC) in the expression of cellular markers **(a)** CD11b, **(b)** F4/80, **(c)** CD3, **(d)** CD4, and **(e)** CD8 was evaluated using qRT-PCR and compared to the housekeeping gene HPRT. Data are shown as scatter plots and mean ± SEM utilizing two-tailed Welch’s *t*-test for analyses. n = 11 for juvenile group (4 female, 7 male). n = 20 for adult group (10 female, 10 male).

**Figure 6.**
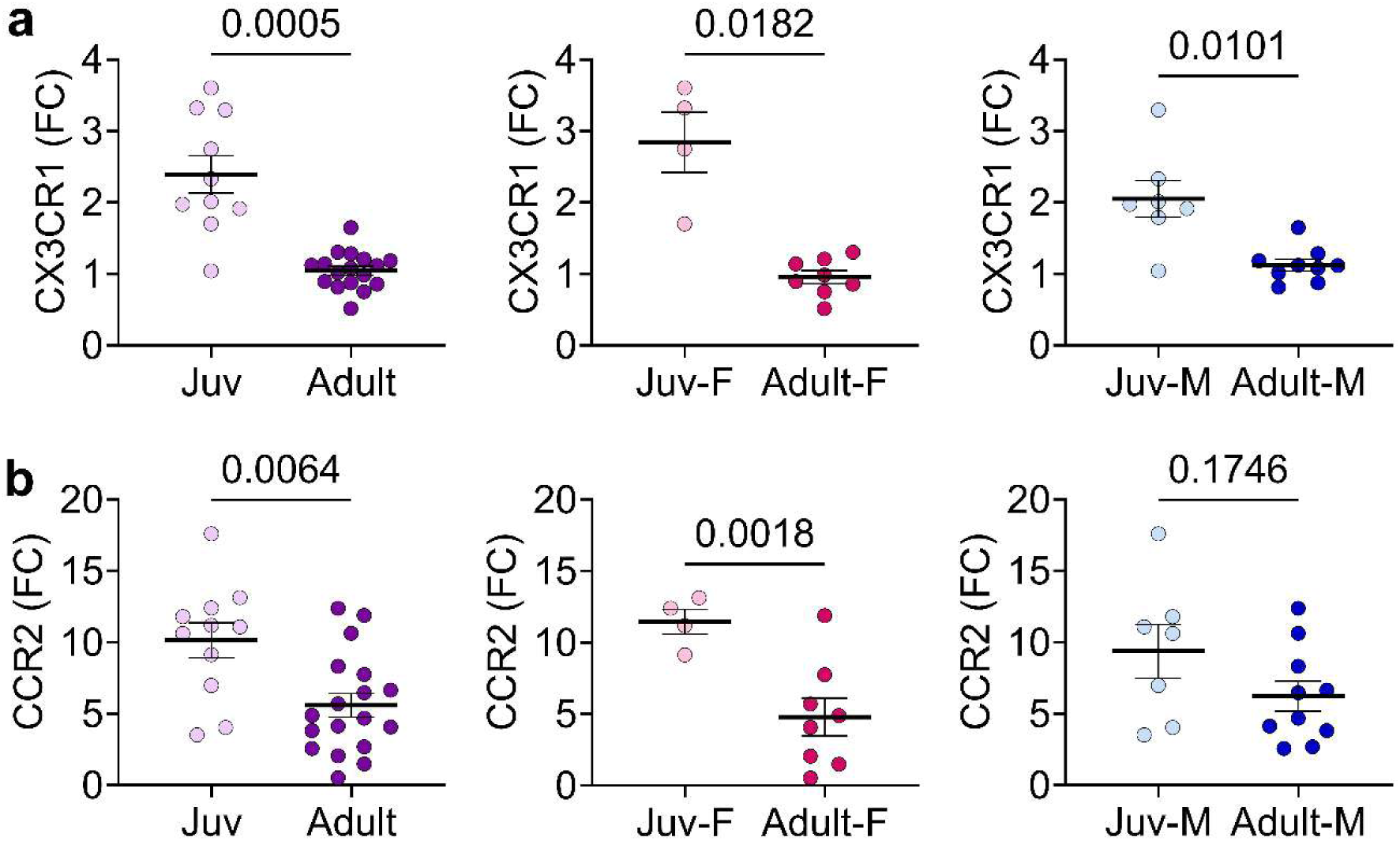
Juvenile female and male mice with myocarditis show differential increases in resident and inflammatory macrophage chemokine receptor marker expression compared with adults with myocarditis. Juvenile and adult female and male mice were injected intraperitoneally with 10^3^ PFU of CVB3 and harvested at day 10 pi. RNA was isolated and used to produce cDNA. Combined groups of juvenile (light purple) and adult (purple) with myocarditis were analyzed. Further sex-separated analyses were conducted between juvenile and adult females (juvenile in pink, adult in red) and males (juvenile in light blue, adult in dark blue) for all targets. Fold change (FC) in the expression of cellular markers **(a)** CX3CR1, and **(b)** CCR2 was evaluated using qRT-PCR and compared to the housekeeping gene HPRT. Data are shown as scatter plots and mean ± SEM utilizing two-tailed Welch’s *t*-test for analyses. n = 11 for juvenile group (4 female, 7 male). n = 20 for adult group (10 female, 10 male).

### Adult, but not juvenile, male mice with myocarditis have lower testosterone levels than healthy controls

In our evaluation of 17β-estradiol (estradiol) and testosterone levels across groups, we found that serum estradiol levels did not differ with age or disease (**Figure 7a**). Similarly, testosterone levels in females were low, as expected, and did not vary with age or due to myocarditis (**Figure 7b, left**). Among males, testosterone showed different trends. Two- way ANOVA showed that the interaction between age and disease was not significant, F (1, 39) = 1.872, p = 0.18, η^2^ = .03. However, there were significant main effects of age, F (1, 39) = 7.637, p = 0.0087, η^2^ = .13, and infection, F (1, 39) = 6.501, p = 0148, η^2^ = .11. Post-hoc testing showed healthy adult males, as expected, had higher serum testosterone levels than healthy juvenile males (p = 0.0118) (**Figure 7b, right**). However, adult males with myocarditis had significantly lower testosterone levels compared with healthy adult males (p = 0.0183) **(Figure 7b, right)**.

**Figure 7.**
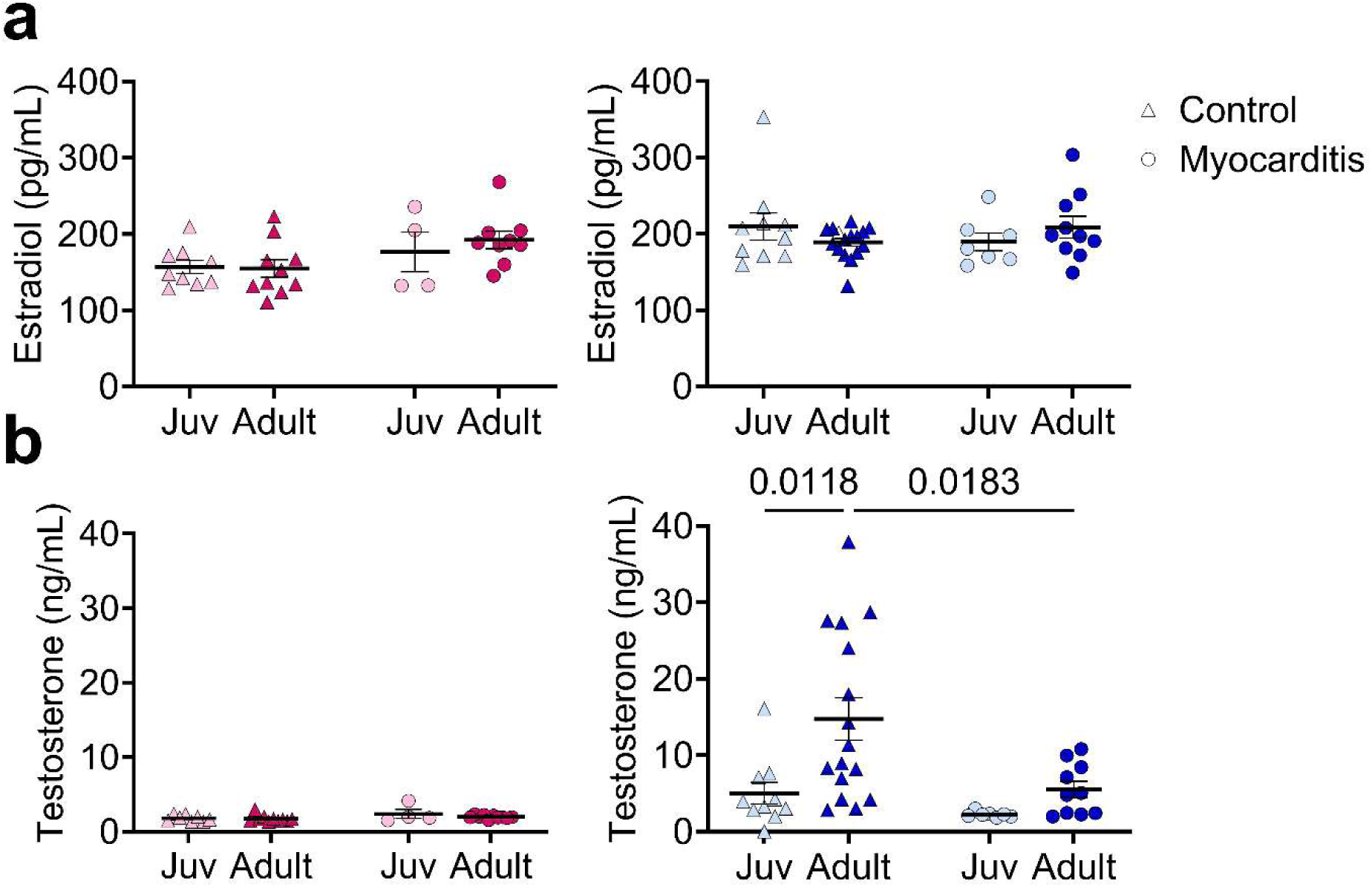
Circulating testosterone, but not 17β-estradiol, is increased in adult vs. juvenile male mice but lower during myocarditis. Juvenile and adult female and male mice were injected intraperitoneally (ip) with 10^3^ PFU of CVB3, and sera were isolated via cardiac puncture at day 10 pi. 17β-estradiol levels and testosterone levels were determined by ELISA and compared between myocarditis (circles) and control groups (triangles). Juvenile females are represented in light pink, juvenile males in light blue, adult females in red, and adult males in dark blue. Data are shown as scatter plots and mean ± SEM utilizing two-way ANOVA with Tukey’s multiple comparisons test. n = 11 for juvenile group (4 female, 7 male). n = 20 for adult group (10 female, 10 male).

### Expression of complement cascade components is increased in both juvenile and adult mice with myocarditis

Given the importance of the complement cascade to innate immunity, we analyzed the expression of its components during CVB3 myocarditis. As expected, across all groups, expression increased in response to infection (data not shown). Expression levels for C3, C4, and C5aR1 were not significantly different between juvenile and adult mice with myocarditis **(Figure 8a, 8b, and 8d)**. Juvenile mice with myocarditis showed significantly higher expression of C3aR1 than adult mice (p = 0.0343) **(Figure 8c)**. By sex, no differences were seen in marker expression.

**Figure 8.**
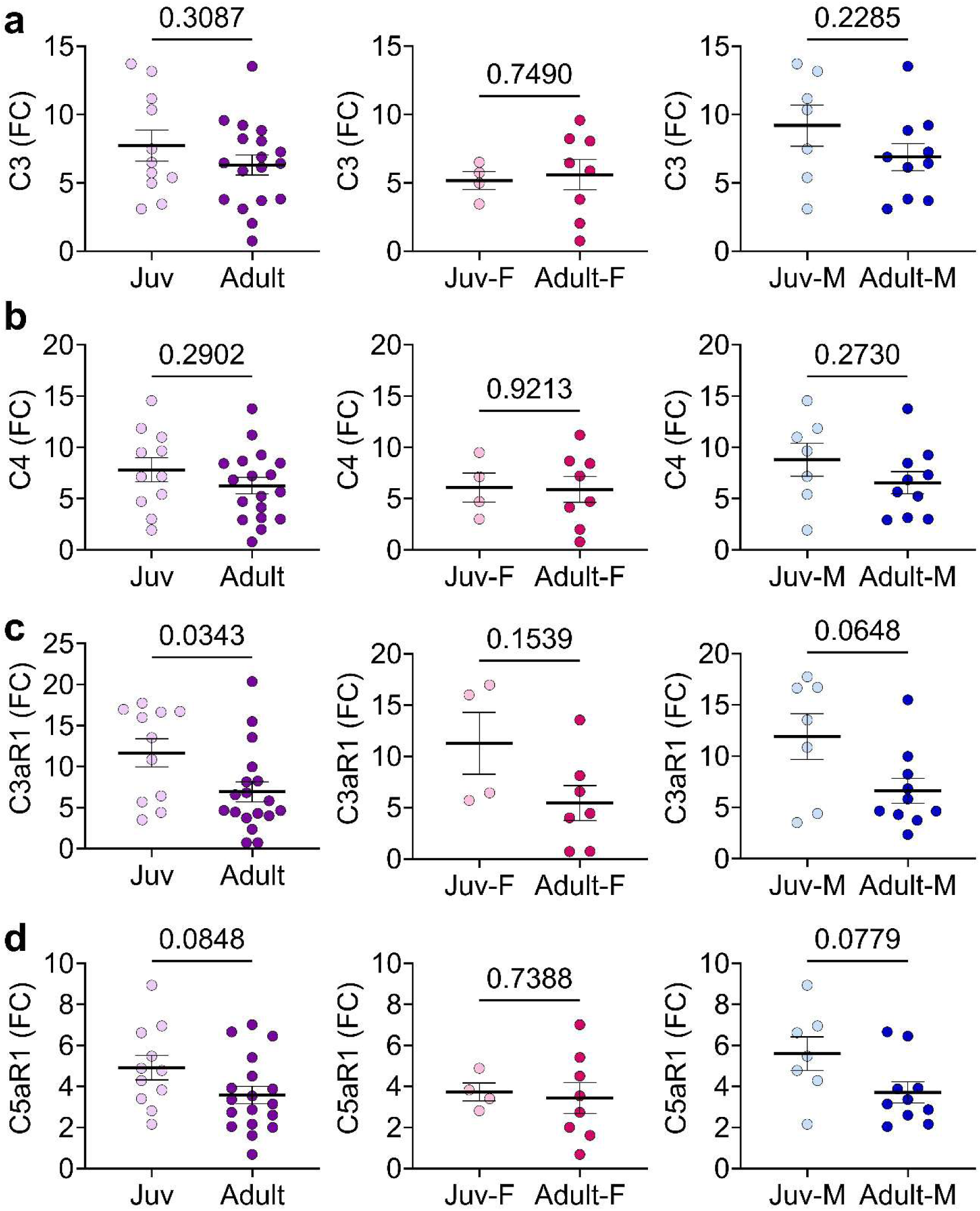
Juvenile mice with myocarditis show increased expression of complement genes compared with adult mice. Juvenile and adult female and male mice were injected intraperitoneally with 10^3^ PFU of CVB3 and harvested at day 10 pi. RNA from juvenile and adult male and female hearts was isolated and used to produce cDNA. Combined groups of juvenile (light purple) and adult (purple) with myocarditis were analyzed. Further sex-separated analyses were conducted between juvenile and adult females (juvenile in pink, adult in red) and males (juvenile in light blue, adult in dark blue) for all targets. Fold change (FC) in the expression of complement markers **(a)** C3, **(b)** C4, **(c)** C3aR1, and **(d)** C5aR1 was evaluated using qRT-PCR and compared to the housekeeping gene HPRT. Data are shown as scatter plots and mean ± SEM utilizing two-tailed Welch’s *t*-test for analyses. n = 11 for juvenile group (4 female, 7 male). n = 20 for adult group (10 female, 10 male).

### Expression of TLRs and the NLRP3 inflammasome show differential increases in juveniles compared with adults

Finally, we investigated changes in TLR and NLRP3 inflammasome expression given their critical role in the viral immune response. Compared with adults with myocarditis, juvenile mice with myocarditis showed significantly increased expression of signaling components TLR2 (p = 0.0282) **(Figure 9a)** and TLR7 (p = 0.0415) **(Figure 9b)** with no difference in TLR4 (p = 0.49) (**Figure 9b)** at this timepoint. Among the terminal components of the pathway, juvenile mice with myocarditis showed increased expression of NLRP3 (p = 0.0068) and caspase-1 (p = 0.0106), but not IL-1β (p = 0.0805) by qRT- PCR, compared with adults with myocarditis **(Figure 9d, 9e, and 9f)**. Separating by sex, neither juvenile males nor females with myocarditis showed increased expression of inflammasome components compared with adults with myocarditis.

**Figure 9.**
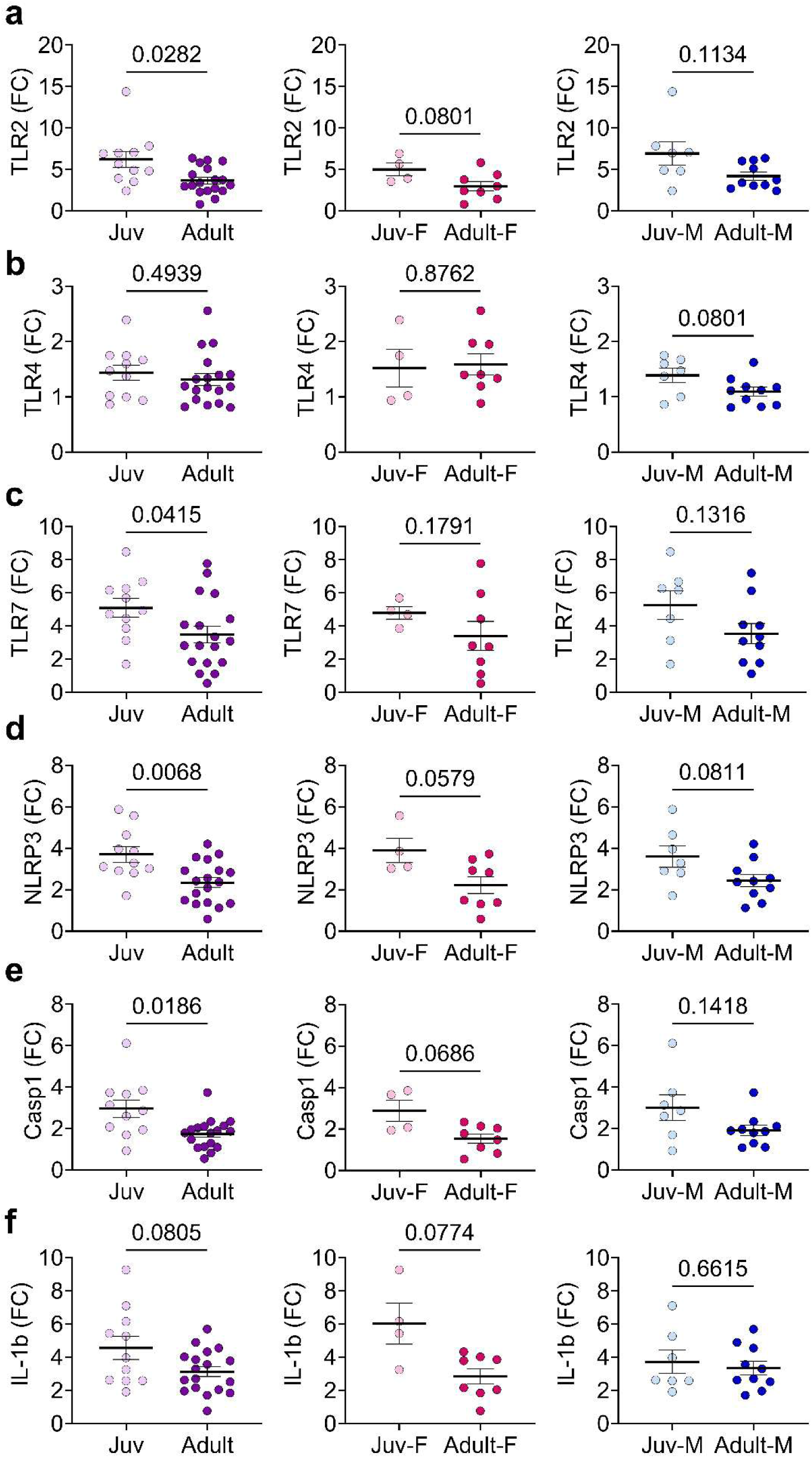
Expression of NLRP3 inflammasome components is increased in juvenile mice with myocarditis compared with adult mice with myocarditis. Juvenile and adult female and male mice were injected intraperitoneally with 10^3^ PFU of CVB3 and harvested at day 10 pi. RNA from juvenile and adult male and female hearts was isolated and used to produce cDNA. Combined groups of juvenile (light purple) and adult (purple) with myocarditis were analyzed. Further sex-separated analyses were conducted between juvenile and adult females (juvenile in pink, adult in red) and males (juvenile in light blue, adult in dark blue) for all targets. Fold change (FC) in the expression of NLRP3 inflammasome components **(a)** TLR2, **(b)** TLR4, **(c)** TLR7, **(d)** NLRP3, **(e)** Casp1, and **(f)** pro-IL-1β was evaluated using qRT-PCR and compared to the housekeeping gene HPRT. Data are shown as scatter plots and mean ± SEM utilizing two-tailed Welch’s *t*-test for analyses. n = 11 for juvenile group (4 female, 7 male). n = 20 for adult group (10 female, 10 male).

## Discussion

In this study, we report the first description of a translational mouse model of juvenile coxsackievirus B3 myocarditis, compared with the established adult model from the Fairweather group.^23^ Overall, we observed that 3-4-week-old mice exhibited more severe cardiac inflammation and necrosis than sex-matched adult mice. Importantly, we also did not observe sex differences in cardiac inflammation in juvenile mice as observed in adults^3,39^ replicating findings in clinical populations. These findings suggest that age plays a critical role in the overall immune response in viral myocarditis and that models specific to the juvenile population are necessary to study disease pathogenesis in this population.

To date, studies of myocarditis in animal models have primarily used mice older than 5-8 weeks.^29,40,41^ These models have identified and investigated sex differences in the adult population and reinforced the role of sex hormones in disease pathogenesis.^42^ Additionally, clinical studies and trials in patients with myocarditis have focused on adult populations.^9,21^ Therefore, extrapolation of the findings of these studies to juvenile, prepubescent populations should be done with caution. There remains a lack of studies utilizing juvenile animals to study viral myocarditis. The only example that attempted to develop a juvenile viral myocarditis mouse model did not result in cardiac inflammation but instead led to neuroinflammation and paralysis.^43^ However, juvenile animal models for other diseases, such as Pertussis, Respiratory Syncytial Virus, and Influenza (although not specifically investigating myocarditis), exist and demonstrate that the immune response is dynamic across the lifespan.^44–46^ Furthermore, studies comparing the immune response in juvenile and adult mice have been conducted. In a model of MRSA pneumonia, infection in neonatal mice resulted in increased bacterial burden and decreased macrophage and neutrophil recruitment compared with infected adult mice.^47^ In the context of the viral immune response, a model of paramyxovirus infection showed that neonatal mice had lower inflammation, decreased cytokine release, and delayed activation of pattern recognition receptors compared with adults.^48^ In Pneumovirus infection, comparisons across ages show that, despite similar viral replication rates, the overall magnitude and type of inflammation vary, with older mice showing the most prominent immune cell recruitment.^49^ Studies using mouse models of influenza have shown that neonatal immune cells tend to have decreased capacity to secrete interferon (IFN)-γ and reduced T cell responses, findings that replicate those in children.^50–53^ Additionally, this decreased response has been associated with reduced viral clearance.^51^

Our model shows both similarities and differences to other investigations of immune responses in juvenile and adult mice. In response to CVB3 infection in adult mice, we observe an expected increase in macrophage infiltration into the heart, as previously reported.^54^ Additionally, our results regarding expression levels of the complement pathway and NLRP3 inflammasome components agree with previously published results in CVB3 infection.^55,56^ However, these processes have never been explored in a juvenile myocarditis model. Our model not only shows that myocardial inflammation is worse in juvenile mice but also that this translates into differences in immune cell infiltration, immune cell marker expression, and innate immunity pathways. Overall, this reinforces the need to characterize this response in juvenile mice and to understand better the differential severity of disease in the juvenile population.

In clinical populations, most pediatric myocarditis research focuses on outcomes. However, some comparative studies have been published. Mainly, it has been shown that, compared with pediatric patients hospitalized with dilated cardiomyopathy, those hospitalized with acute myocarditis exhibit increased expression of terminal NLRP3 inflammasome components at both the gene and protein levels.^57^ However, this study did not include adult patients with myocarditis or control patients. As expected, our model showed increased expression of inflammasome components in the myocarditis groups compared with the control group and revealed differences between pediatric and adult myocarditis groups.

One limitation of our study is that our ability to precisely age the mice was limited by the method we used to acquire them. While puberty checks allowed us to estimate age based on puberty status, future experiments using timed pregnancies would provide even more accurate ages. Additionally, although we report differences by sex, these results do not confirm true sex differences by age, as our sample size, particularly for juvenile females, is small. Thus, while our comparisons of juvenile and adult groups regardless of sex are appropriately powered, further investigation through additional experiments is needed to establish the validity of our preliminary findings of age differences within each sex. Another limitation of this study is that we are assessing only one time point; future directions should investigate sex and age differences at earlier time points during myocarditis development and progression to dilated cardiomyopathy using our translational animal model of viral myocarditis. Finally, our model utilized the same viral dose across all groups. This may lead to differences in relative dose between pediatric and adult mice due to the inherent differences in size. It is possible that this could explain the differences seen. Future studies will explore the use of a viral dose normalized to body weight to better control for potential dosing inconsistencies.

### Perspectives and Significance

Overall, myocarditis continues to present a significant burden of disease. Treatment is primarily focused on symptomatic relief of associated heart failure rather than treatment of the disease itself. Moreover, treatment strategies in pediatric populations are derived from that of adult populations.^17^ However, it has been well established that the immune response in children differs from that in adults. Despite this, no studies have compared responses in these two populations. Here, we present the first report of an animal model designed to study mechanisms of CVB3 myocarditis and how they differ between pediatric and adult populations. We show that, compared with adult mice with myocarditis, juvenile mice exhibit greater inflammation and potential sex-dependent differences in immune pathway marker expression. Ultimately, our findings highlight key differences between these groups and reinforce the need for ongoing studies to identify new individualized targets for disease-specific diagnostics and therapies.

## Declarations

### Ethics Approval and Consent to Participate

Mice were used in strict accordance with the recommendations in the Guide for the Care and Use of Laboratory Animals from the National Institutes of Health. Approval was obtained from the Animal Care and Use Committee at Mayo Clinic, Florida. Mice were sacrificed according to the Guide for the Care and Use of Laboratory Animals from the National Institutes of Health.

### Consent for publication

This manuscript does not contain patient data requiring consent for publication.

### Availability of data and materials

All data used and/or analyzed during the current study are available from the corresponding author upon reasonable request.

### Conflict of Interest Disclosure

LTC is a consultant for Moderna, Cardiol Therapeutics and Foresee Pharmaceuticals. LTC is an Equity Owner in Stromal Therapeutics. All other authors declare no conflicts of interest.

### Funding

This work is partially supported by the NIH National Heart Lung and Blood Institute (NHLBI) under award number R01 HL164520 (DF), National Institute of Allergy and Infectious Diseases under award numbers R21 AI180863-01A1 (KAB), R21 AI163302 (KAB), the American Heart Association under award number 23SCEFIA1153413 (KAB), the Mayo Clinic Department of Immunology (KAB), the For Elyse Foundation (KAB), the Myocarditis Foundation (KAB), and the Mayo Foundation for Medical Education and Research (LTC). The content is solely the responsibility of the authors and does not necessarily represent the official views of the funding agencies.

### Authors’ contributions

Conceptualization: KB. Funding acquisition: DF, KB. Sample acquisition: LB, EW, DB, MS, DD, SK, DF, KB. Methodology: JR, LM, EW, KG, CD, KR, PT, AD, NF, LP, DB, MS, DD, DF, SK, KB. Data Analysis: JR, LM, EW, KG, DB, DF, KB. Data Curation: JR, DG, KB. Project Administration: DF, KB. Writing original draft: JR, LTC, JP, DF, DG, KB. Writing, reviewing, and editing: JR, LM, EW, KG, CD, KR, PT, AD, NF, LP, DB, MS, DD, SK, JNE, LTC, JP, DF, DG, KB.

## Acknowledgements

The authors would like to express their appreciation to the For Elyse Foundation and the Myocarditis Foundation for providing funding for this study. We thank the Dickson Histology Group for their work in embedding and staining slides for this project. This group includes Dr. Dennis W. Dickson, Linda Rousseau, Virginia Phillips, Ariston Libraro, and Monica Castanedes. We would also like to acknowledge the University of Florida Molecular Pathology Core and Mayo Clinic for their assistance in generating tissue sections for picrosirius red staining.

